# ChromVAR: Inferring transcription factor variation from single-cell epigenomic data

**DOI:** 10.1101/110346

**Authors:** Alicia N. Schep, Beijing Wu, Jason D. Buenrostro, William J. Greenleaf

## Abstract

Single cell ATAC-seq (scATAC) yields sparse data that makes application of conventional computational approaches for data analysis challenging or impossible. We developed chromVAR, an R package for analyzing sparse chromatin accessibility data by estimating the gain or loss of accessibility within sets of peaks sharing the same motif or annotation while controlling for known technical biases. chromVAR enables accurate clustering of scATAC-seq profiles and enables characterization of known, or the *de novo* identification of novel, sequence motifs associated with variation in chromatin accessibility across single cells or other sparse epigenomic data sets.

## Main Text

Transcription factor binding to regulatory DNA sequences controls the activity of *cis*-regulatory elements, which modulate gene expression programs that define cell phenotype. Assays for probing chromatin accessibility have enabled the discovery of *cis*-regulatory elements and *trans*-acting factors across different cell states and types that lead to functional changes in gene expression^1^. Concurrently, single cell genomic and transcriptomic methods have enabled unbiased *de novo* deconvolution of dynamic or diverse cellular populations^2,3^. Recently developed assays for measuring chromatin accessibility within single cells^4,5^ promise to enable the identification of causative *cis-* and *trans-* regulators that bring about these diverse cellular phenotypes.

However, the exceedingly sparse nature of single cell epigenomic data sets present unique and significant computational challenges. All single-cell epigenomic methods are intrinsically sparse, as the total potential signal at a genomic locus is fundamentally limited by the copy number of DNA, thus generating 0, 1 or 2 reads from regulatory elements within a diploid genome. Methods developed for single cell RNA-seq have shown that measuring the dispersion of gene sets, such as Gene Ontology or co-expression modules, rather than individual genes can be a powerful approach for analyzing sparse data^6^. In this vein, and building on previous work^4,7^, we have developed *chromVAR*, a versatile R package for analyzing sparse chromatin accessibility data by measuring the gain or loss of chromatin accessibility within sets of genomic features while controlling for known technical biases in epigenomic data. We show that *chromVAR* can be used to identify transcription factor (TF) motifs that define different cell types and vary within populations, providing a unique analytical toolkit for analysis of sparse epigenomic data.

The chromVAR package takes as inputs 1) aligned sequencing reads, 2) chromatin accessibility peaks (derived from either data aggregated across cells or external resources), and 3) a set of chromatin features representing either motif position weight matrices (PWMs) or genomic annotations (Figure 1A, Figure S1). For use as input (3) into chromVAR, we have curated a set of human and mouse PWMs from the cisBP database^8^ that represent a diverse and comprehensive collection of known TF motifs. Alternately, user provided TF motifs or other types of genomic annotations, such as enhancer modules, ChIP-seq peaks, or GWAS disease annotation may be used. chromVAR may also be applied to a collection of kmers--DNA sequences of a specific length k-- in order to perform an unbiased analysis of DNA sequence features that correlate with chromatin accessibility variation across the cells or samples.

**Figure 1.**
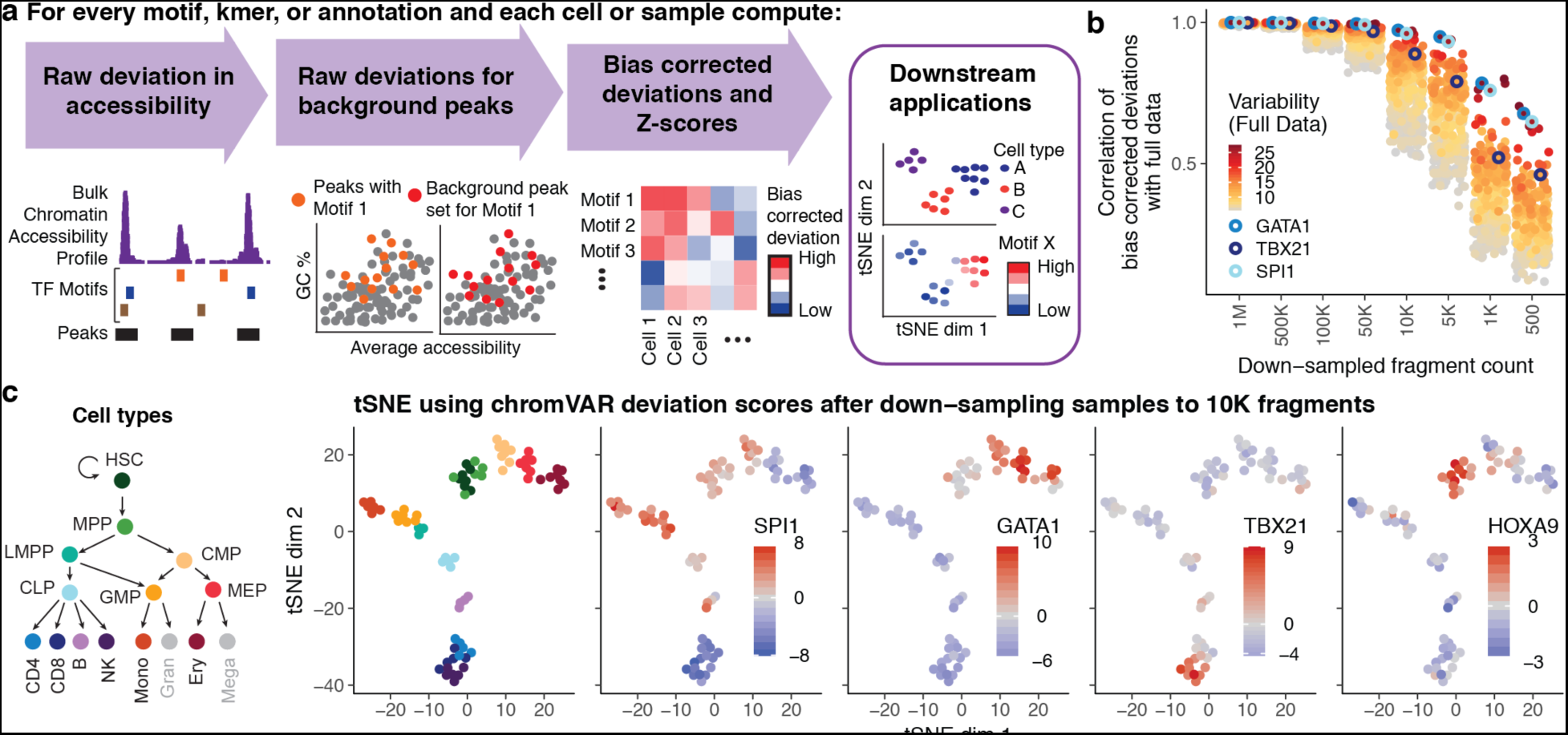
*chromVAR* enables interpretable analysis of sparse chromatin accessibility data. a) Schematic illustrating how chromVAR uses aggregation of accessibility across peaks sharing a common feature (e.g. a motif) with bias correction to generate scores for each cell or sample that can be used for downstream analysis b) Correlation of chromatin accessibility bias corrected for 77 samples from different hematopoietic populations before and after down-sampling total sequencing reads from full data. Each point shows the correlation for a different motif. The top 20 % most variable motifs are shown. Three of the most variable motifs are highlighted. c) tSNE visualization of different samples using normalized deviations calculated from data down-sampled to 10,000 fragments per sample. In the first panel, cells are colored by cell type, and in other panels cells are colored by the deviation scores for different motifs.

chromVAR first computes a “raw accessibility deviation” for each motif and cell, representing the difference between the total number of fragments mapping to peaks containing the given motif and the total expected number of fragments based on the average of all input cells. Technical biases between cells due to PCR amplification or variable Tn5 tagmentation conditions can lead to differences in the number of observed fragment counts between cells for a given peak set with distinct GC content or mean accessibility (Figures S2). To account for these technical confounders, “background” peak sets are created for each annotation, which comprise of an equal number of peaks matched for GC content and average accessibility (Figure S3–Figure S6). The raw accessibility deviations for these background peak sets are used to compute a bias corrected deviation and Z-score for each annotation and cell, providing a bias-corrected differential measure describing the gain or loss of accessibility of a given genomic annotation relative to the average cell profile (see methods). These bias corrected deviations and Z-scores can be used for a number of downstream analyses, including *de novo* clustering of cells and identification of key regulators that vary within and between different cell types. The chromVAR package includes a collection of tools for such downstream analysis, including an interactive web application for exploring the relationship between key TF motifs and clustering of cells (Figure S7). We have also incorporated tools for generating previously-described analyses characterizing the correlation and potential cooperativity between two TF binding sites within the same regulatory element, and computing chromatin variability across regions in *cis*^4^.

To test the applicability of this computational workflow for single-cell analysis, we set out to measure the robustness of chromVAR outputs to data downsampling. To do this, we applied chromVAR to bulk ATAC-seq data from a deeply-sequenced set of hematopoietic cell types^7^, and compared the results of the analysis for the data across various degrees of downsampling. We found that the TF motif deviations using 10^6^ to 5x10^3^ fragments per sample are highly correlated to those determined using the full data set (Figure 1B, S8). The clustering accuracy using the bias corrected deviations is also largely preserved after downsampling, and compares favorably to clustering using PCA (Figure S8; see methods).

Importantly, chromVAR provides robust results for 10,000 fragments per cell, a typical number of fragments generated from a single-cell using scATAC-seq^4^. By projecting the vector of bias corrected deviations from individual cells into two dimensions using tSNE^9^, *chromVAR* enables the reconstruction of the major hematopoietic lineages using 10,000 fragments per sample. With this analytical framework, we can also visualize the TFs associated with significant chromatin accessibility within each simulated single cell epigenome, thereby correctly identifying known master regulators of hematopoiesis, including HOXA9, SPI1, TBX21, and GATA1^10–13^ (Figure 1C).

We next characterized chromVAR’s ability to capture biologically relevant chromatin variability from single cell ATAC-seq data drawn from multiple distinct cell lines and human samples (Figure S9). Using tSNE with bias corrected deviations for motifs and 7mers, we clustered individual cells into distinct cell types and observe individual motifs that best distinguish each cell type (Figure 2A). Notably, well-defined, distinct clusters are formed in this tSNE projection when using the bias corrected deviations, but the clustering is confounded by technical biases when using raw deviations without the bias correction infrastructure. Interestingly, we also observe that cells from acute myeloid leukemia (AML) patients cluster between lymphoid-primed multipotent progenitors (LMPPs), monocytes, and HL60 (an AML derived cancer cell line) cells. In this unsupervised analysis, we find that the AML leukemic stem cells are more similar to LMPPs, while the AML blasts are more similar to the monocytes. In addition, we also observe that patient 1 (AML blast 1) maintains a more stem-like state when compared to patient 2 (AML blast 2) as anticipated from alternate analyses of these cells^14^. By visualizing the cell-specific Z-scores layered on this projection, we identify putative TFs that may promote the stem-like versus differentiated leukemia phenotype; for example, the master-regulators of myeloid cell development SPI1 (PU.1) and CEBPA^15^ appear as the most differential motifs between AML leukemic stem cells (LSCs) and blasts (Figure 2b-d).

**Figure 2.**
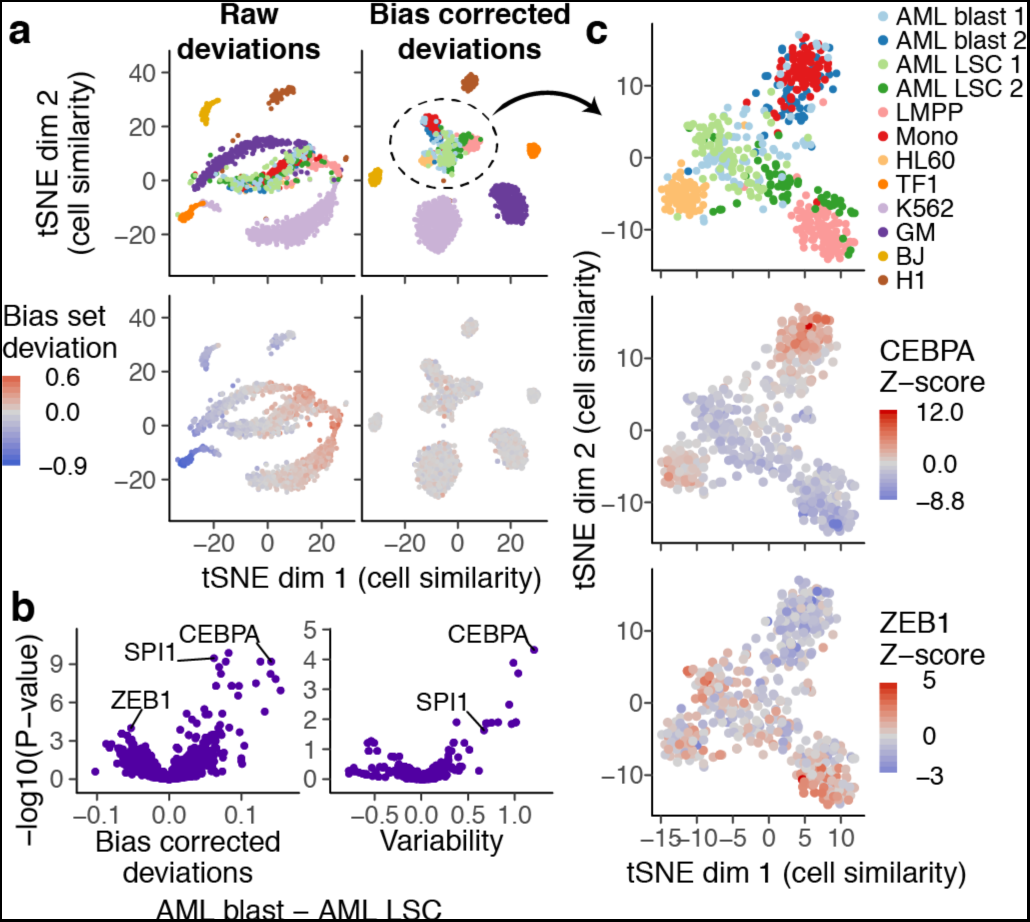
*chromVAR* enables clustering of single cell populations and interpretation of motifs underlying chromatin accessibility variation in single cells. a) tSNE visualization of similarity of single cells based on chromVAR raw (left) or bias corrected deviations (right) for motifs and 7mers (see methods). In top panels, points are colored by cell type and in bottom panels points are colored by raw (left) or bias corrected (right) calculated deviations for a set of random peaks with high GC content and high average accessibility (the bias set). b) Volcano plot showing the mean difference in bias corrected accessibility deviations (left) and variability (right) for each motif between the AML blast and LSC cells versus the −log10(P-value) for that difference. c) tSNE with bias corrected deviations for AML blast and LSC, monocyte, LMPP, and HL60 cells. In top panel, points are colored by cell type, and in other panels points are colored by deviation Z-scores for CEBPA and ZEB1 respectively.

In addition to visualizing the similarity of cells, we inverted our tSNE analysis to visualize the similarity of motifs and kmers in their activity patterns across cells (Figure 3a). In this visualization, motifs and kmers that have similar activity profiles across cells cluster together in the tSNE subspace, allowing the identification of major clusters representing several different TF families. Notably, different TFs within the same family (e.g. GATA1 and GATA2) often bind highly similar motifs, and therefore chromVAR alone cannot distinguish the causative regulator binding a particular TF motif. In the inverted tSNE visualization for motif and kmer similarity, most, but not all kmers cluster with a known motif, suggesting k-mer analysis may enable *denovo* discovery of previously unannotated motifs.

**Figure 3.**
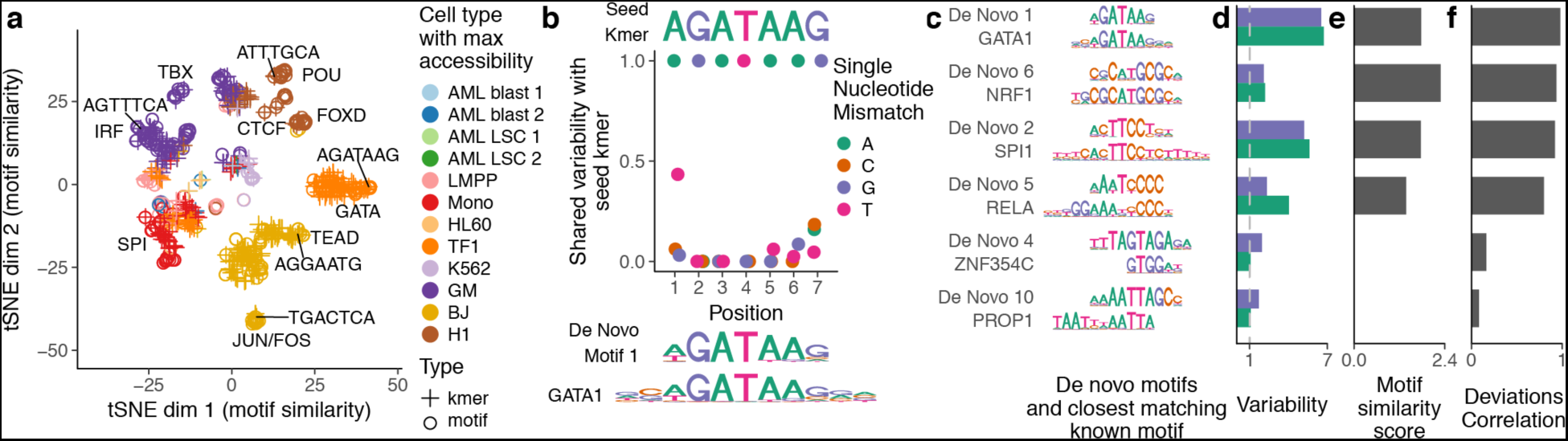
*chromVAR* identifies *de novo* motifs associated with chromatin accessibility variation in single cells. a) tSNE visualization of similarity between motifs and kmers based on the vector of normalized deviations across different cells. Labels highlight predominant families of motifs within a cluster and example 7mers b) For the seed kmer “AGATAAG”, the shared variability of k-mers with one mismatch from the seed kmer. These shared variabilities can be used to assemble a *de novo* motif that closely resemble the GATA1 motif (see methods). c) Example *de novo* motifs assembled by chromVAR using deviation scores for 7-mers, along with the closest matching known motif below it. d) Variability for both the *de novo* motif and the known motif for each pair in panel (c). e) Motif similarity score (see methods) between the *denovo* motif and the known motifs in (c) f) The correlation between the normalized deviations of the *de novo* motif and the known motif for each pair in (c).

By comparing the variation in chromatin accessibility across cells between highly similar kmers, we can identify critical bases associated with chromatin accessibility variation. For example, the “AGATAAG” kmer, which closely matches the GATA1 motif, is highly variable across single cells, but most kmers differing by one nucleotide share little or none of that variability (Figures 3b, S10). The mismatched kmer with the greatest correlated variability is “TGATAAG”, which is consistent with the weights of each nucleotide in the GATA1 motif.

We can use these comparisons of variation between highly similar kmers to construct *denovo* motifs representing sequences associated with variation in chromatin accessibility. In brief, we start with highly variable “seed” kmers, and use the covariance between the seed kmer and kmers either differing by one mismatch or partially overlapping the seed kmer to assign weights to different nucleotide bases at each position of the motif model (see methods). Importantly, many *de novo* motifs assembled using this approach closely match known motifs (Figure 3c-f, S11). For motifs that do not closely match to a known TF, we confirmed that the constructed motifs were also associated with variation in DNase hypersensitivity between different samples represented in the Roadmap Epigenomics Project^16^ (Figure S12), demonstrating that these *denovo* motifs are associated with chromatin accessibility variation in two distinct accessibility assays. To further validate the discovery of these putative *trans*-regulators we calculated aggregate TF “footprints”, a measure of the DNase or Tn5 cut density around the given motif, and found a diverse set of accessibility profiles (Figure S13). Interestingly, several of these motifs did not match canonical narrow (~20 bp) transcription factor footprints, but rather are associated with a large footprint (>20 bp) potentially indicative of larger regulatory complexes.

In summary, we envision that *chromVAR* will be broadly applicable to single-cell and bulk epigenomics data to provide an unbiased characterization of cell types and the *trans* regulators that define them. As methods for measuring the epigenome in single-cells and bulk populations continue to improve in throughput and in quality, scalable analytical infrastructure is needed. Analysis workflows for ATAC or DNase-seq data often include the identification of motifs enriched in differentially accessible peaks, but such approaches scale poorly to comparisons across many sample types and require sufficient read depth per-locus to determine differential peak accessibility. In contrast, *chromVAR* analysis is highly robust to low sequencing depth and readily scales to hundreds or thousands of cells or samples. Budget-constrained researchers often face a trade-off between the number of samples to sequence and the sequencing depth for each sample; sparse sequencing analysis coupled with chromVAR analysis may enable new applications of “bulk” ATAC, DNase-seq or other epigenomic methods as large-scale screening tools. We also anticipate that *chromVAR* will enable additional downstream analyses of single cell chromatin accessibility data, as the reduction of dimensionality associated with vectors of bias corrected deviations provide a powerful input to existing algorithms for inferring spatial and temporal relationships between cells.

## Methods

### ATAC-seq, scATAC, and DNase Data

In addition to the previously published data, we generated three new replicates of single-cell K562s using the previously published protocol^4,7^. Bulk ATAC-seq and scATAC-seq data was aligned and filtered as described previously^4,7^. Uniformly processed DNase data was downloaded from the Roadmap Epigenomics Project Portal^16^.

### Peaks

For the bulk data analysis, DNase hypersensitivity peaks from the Roadmap Epigenomics Project were used. MACS2^17^ peaks for blood cells (Primary monocytes from peripheral blood, Primary B cells from peripheral blood, Primary T cells from peripheral blood, Primary Natural Killer cells from peripheral blood, Primary hematopoietic stem cells G-CSF-mobilized Female, Primary hematopoietic stem cells G-CSF-mobilized Male, and Monocytes-CD14+ RO01746 Cell line) were downloaded from the Epigenomics Roadmap Portal^16^. For the single cell ATAC-seq data, peaks were called for each cell line or type using MACS2 applied to the merged single cell ATAC-seq data. All peaks were re-sized to a uniform width of 500 bp, centered at the summit. For both the set of peak calls from the blood cells in Roadmap and the set of peak calls from the scATAC-seq data, peaks were combined by removing any peaks overlapping with a peak with greater signal.

### Motif collection and motif matching

We curated Position Frequency Matrices from cisBP representing a total of 15,389 human motifs and 14,367 mouse motifs. To filter motifs to a representative subset, we first categorized motifs as high, medium and low quality, as is provided in the cisBP database. We then grouped all 870 unique human or 850 unique mouse TF regulators represented in the database and assign these regulators to their most representative TF motif(s). To do this, we iterated through each regulator to find all TF motifs associated with that regulator from the high-quality motif list. For these associated motifs, we first computed a similarity matrix using the pearson correlation of the motifs. To select a representative subset from this list, we chose the largest group of associated similar motifs (R>0.9), and selected the motif most similar to the mean of the group. Motifs with an R>0.9 to the chosen motif were discarded from further analysis, and the process was iterated until no motifs remained. We repeated the process for regulators not associated to any motif in the high-quality database using the medium and then low-quality databases. The final curated motif database contains 1,764 human motifs and 1,346 mouse motifs representing 870 human and 850 mouse regulators. The resulting names are formatted as follows∷“ensemble ID”_”unique line number”_”common TF name”_”direct (D) or inferred (I)”_”number of similar motifs”. These position frequency matrices were then converted into Position Weight Matrices (PWMs) by taking the log of the frequency after adding a 0.008 pseudocount and dividing by 0.25.

These PWMs were used for all analyses, except for Supplementary Figure S4–Supplementary Figure S6 in which a smaller set of motifs from the JASPAR CORE database 2016 were used^18^.

The MOODS^19^ C++ library (Version 1.9.3) was used for identifying peaks containing a motif match, using a p-value cutoff of 5x10^-5^. As background frequencies we used the nucleotide frequencies across all peaks. We wrapped the MOODS library into an R package, *motifmatchr*, which enables fast determination of motif presence or positions within genomic regions. The package is available at www.github.com/GreenleafLab/motifmatchr.

### Bias corrected deviations and scores

Raw accessibility deviations for a motif or annotation are computed by summing the number of fragments mapping to a peak containing that motif or annotation for a given cell or sample and subtracting the expected number of fragments mapping to the same peaks for that cell or sample, and then dividing the result by the expected number of fragments mapping to the same peaks. The expected number of fragments is determined by computing the fraction of all fragments within peaks mapping to the peaks sharing the motif or annotation across all cells or samples and multiplying that fraction by the number or fragments mapping to peaks in that cell or sample.

For each motif or genomic annotation, background peak sets are sampled that match the set of peaks with the motif or genomic annotation in terms of the distribution of GC content and average accessibility. These background peak sets are determined by finding possible background peaks for each peak using the following procedure: The state space of GC content and the log of the average accessibility of peaks is transformed by the Mahalanobis transformation in order to remove the correlation between the two variables. This transformed space is split into an even grid of bins with a specified number of divisions (50) along each axis evenly spaced between the minimum and maximum values. For a peak in a given bin j, the probability of selecting another peak x in bin i is given by:

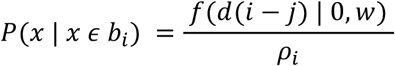

Where f is the probability distribution function of the normal distribution with mean zero and standard deviation w (set to 0.01), *ρ* is the number of peaks in the bin j, and *d*(*i - j*) is the distance between bins i and j.

For each background iteration, raw accessibility deviations were computed for each motif. For all figures except Figure S6, 50 background iterations were used. The bias corrected deviations were computed by subtracting the mean of the background raw deviations for a given motif from its raw deviation. The deviation Z-scores for a motif were computed by dividing the bias corrected deviations by the standard deviation of the background raw deviations for that motif.

### Variability

The variability of a TF motif across samples or cells was determined by computing the standard deviation of the Z-scores. The expected value of this metric is one if the motif peak sets are no more variable than the background peak sets for that motif.

### Downsampling Analysis

To downsample a sample with X total fragments to a depth of Y total fragments, we use the fragment count matrix and for each fragment within a peak retained each fragment with probability X/Y. Thus the downsampled samples are equivalent to having approximately Y total fragments, but not precisely.

The set of peaks used for the analysis remained the same for each down-sampled data set, as the peaks used were from an external data source (Roadmap Epigenomics Project).

For clustering samples using chromVAR results, highly correlated motifs were first removed and then one minus the pearson correlation of the bias corrected deviations was used as the distance matrix for input into hierarchical clustering. For clustering samples using PCA, PCA was performed on the log of the fragment counts for all peaks, and clustering was performed on the euclidean distance between the first five principal components.

### Differential Accessibility and Variability

For determining differentially accessible motifs between AML LSC and blast cells, an unequal variances t-test was used on the bias corrected deviations. For determining differential variability, a Brown–Forsythe test was used on the deviation Z-scores.

### Sample similarity tSNE

For performing sample similarity tSNE, highly correlated motifs or kmers as well as motifs or kmers with variability below a certain threshold (1.5) were first removed from the bias corrected deviations matrix. The transpose of that matrix was then used as input to the Rtsne package^20^, with a perplexity parameter of 8 used for the down-sampled bulk hematopoiesis data and 25 for the single cell ATAC-seq data.

### Motif and kmer similarity tSNE

For performing motif similarity tSNE, motifs or kmers with variability below a certain threshold (1.5) were first removed from the bias corrected deviations matrix, which was then used as input to the Rtsne package ^20^ with perplexity parameter set to 15.

### Kmer Normalized Covariance

As a measure of the shared variability in chromatin accessibility between a reference kmer (or motif) and other kmers (or motifs), we compute a normalized co-variance based on deviation Z-scores. This normalized covariance is simply the covariance of the Z-scores across each cell divided by the variance of the Z-scores for the reference kmer (or motif).

### De novo motif assembly

For assembling *de novo* motifs, we start with the kmer associated with the greatest variability in chromatin accessibility across the cells as a “seed” kmer. We first find the distribution of the normalized covariances between that seed kmer and all other kmers with an edit distance from that seed kmer of at least 3; these values are used as a null distribution for testing the significance of the observed covariances for kmers with a single nucleotide mismatch using a Z-test. For each position along the kmer, the nucleotide of the seed kmer is given a weight of 1. Each alternate nucleotides is given a weight of zero if the p-value for the normalized covariance of the kmer with that mismatch is greater than 0.05; if the p-value is less than 0.05 the nucleotide is given a weight equal to the square of the normalized covariance. The weights for each base pair are then normalized to sum to 1. To further extend the *de novo* motif, we used kmers overlapping the seed kmer with an offset of 1 or 2 bases. For the two bases immediately outside the seed kmer, the weighting of each nucleotide is given by *x ∗ y^2^ + (1 − x)) ∗ 0.25*, where *y^2^* is the square of the normalized covariance for the kmer with the given nucleotide offset (if significant at 0.05 and otherwise 0) and *x* is the maximum value of the normalized covariances for the four kmers (bounded by 0 and 1). For the bases offset by two from the seed kmer, the weighting is computed in the same way except that there are four possible kmers with a given nucleotide at that position that overlap the seed kmer; only the kmer with the maximum normalized covariance with the seed kmer is used.

### Motif Similarity Scores

To score the similarity between a *de novo* motif and the most similar known motif, we first computed the normalized Euclidean distance between the *de novo* motif and all the known motifs in our collection using the optimal local alignment with at least five overlapping bases. We then selected the known motif with the lowest distance as the closest match. The similarity score was computed as the negative of the Z-score for this distance using the distribution of distances for all the motifs in the collection.

### Software Availability

ChromVAR is freely available under the open source MIT license at www.github.com/GreenleafLab/chromVAR.

## Acknowledgments

This work was supported by National Institutes of Health (NIH) P50HG007735 (to W.J.G.), U19AI057266 (to W.J.G.), the Rita Allen Foundation (to W.J.G.) and the Baxter Foundation Faculty Scholar Grant and the Human Frontiers Science Program (to W.J.G). J.D.B. acknowledges support from the Harvard Society of Fellows and Broad Institute Fellowship. ANS acknowledges support from the National Science Foundation (NSF) GRFP (DGE-114747). We thank Caleb Lareau for valuable suggestions for improvements on the package as well as members of Greenleaf and Buenrostro labs for useful discussions.

### Author Contributions

A.N.S, J.D.B, and W.J.G conceived the project and wrote the manuscript. A.N.S. wrote the chromVAR R package and performed the analyses with input from J.D.B and W.J.G. B.W. generated the scATAC-seq data.

---

**Figure S1.**
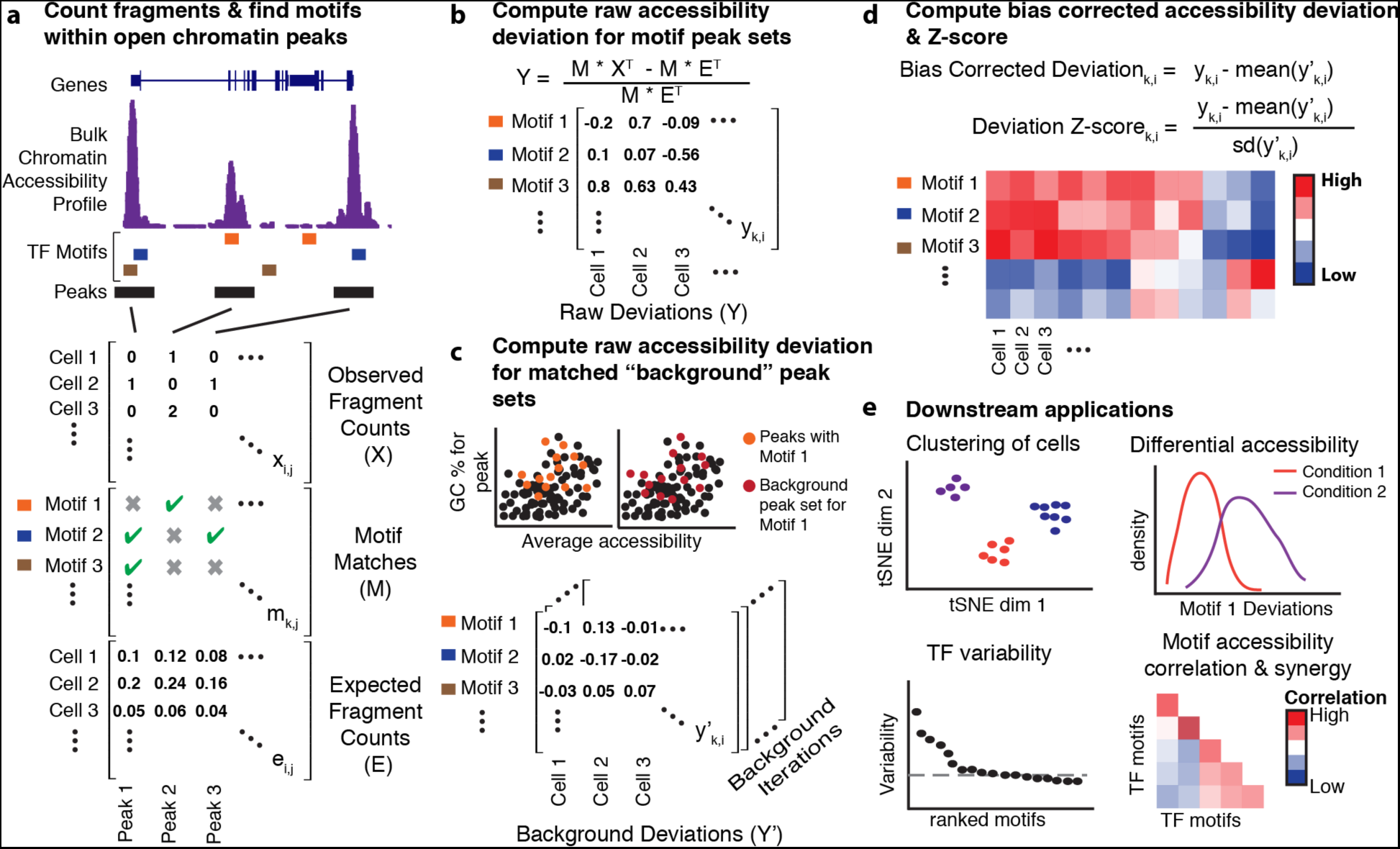
*chromVAR* workflow. a) First, 1) the number of fragments per peak is determined for each cell, then 2) motifs or annotations of interest are assigned to peaks, and 3) the expected fragment count per peak per cell is determined assuming identical read probability per peak for each cell with a sequencing depth matched to that cell’s observed sequencing depth.b) A “raw deviation” is calculated for each motif or annotation feature by summing the fragments in all peaks that contain that feature, then subtracting then dividing by the expected number of fragments in all peaks containing that feature. c) A “raw deviation” is computed for “background sets” of peaks matched in GC content and fragment count to the sets of peaks containing the features of interest. d) The raw deviations for the background sets are used to compute a bias corrected deviation and deviation Z-score for each feature. e) Bias corrected deviations and Zscores
can be used for a variety of downstream applications.

**Figure S2.**
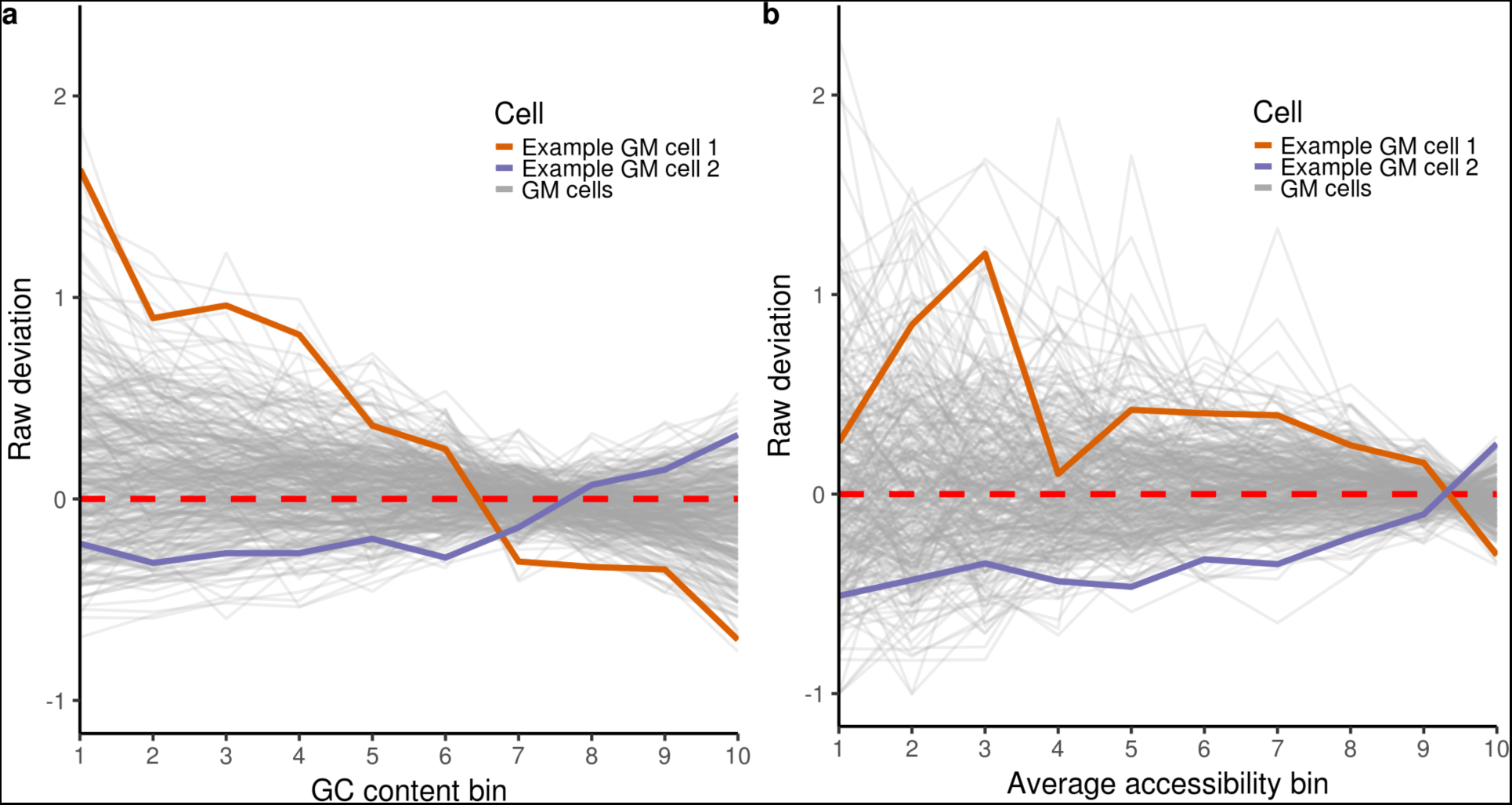
Raw deviations for peak sets defined by GC content or average accessibility. a) Peaks were grouped into 10 bins based on GC content from lowest to highest. The raw deviation in accessibility for each bin was computed for GM cells. Two particular cells are highlighted in orange and purple. b) Same as (a) except bins defined by average accessibility across the cells.

**Figure S3.**
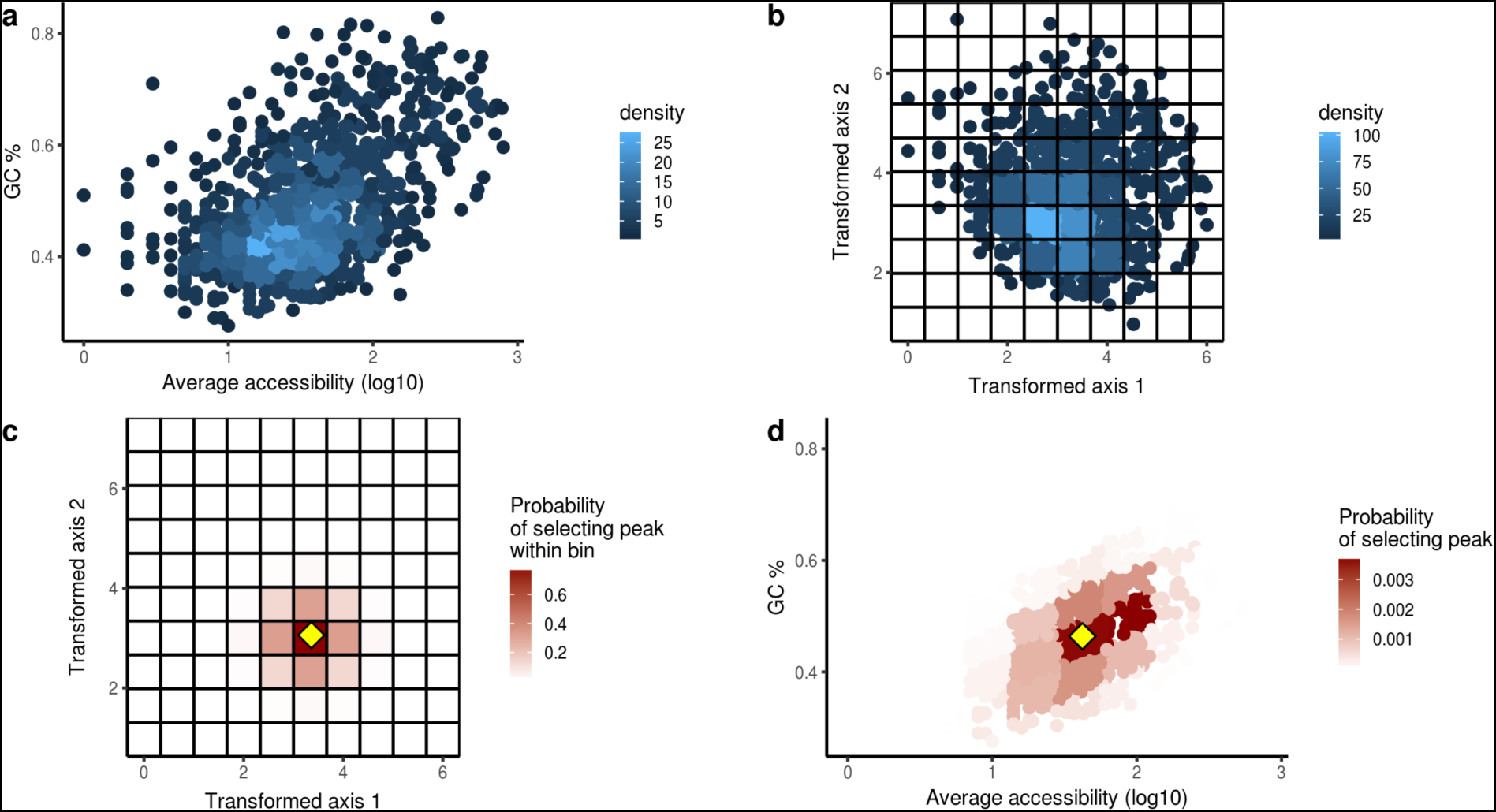
Background peak set selection example. For each background iteration, each peak is assigned a background peak that is similar in GC content and average accessibility. a) GC content versus average accessibility for peaks. b) Same data from (a) after Mahalanobis transformation. Peaks are placed into “bins” based on the values of this transformed data; the grid lines show the boundaries of these bins. c) For the peak indicated with the yellow diamond, the probability of selecting a peak from within a given bin as its background peak. d) Similar to (c), but showing the probability of selecting an individual peak as the background peak for the indicated peak in terms of the untransformed space.

**Figure S4.**
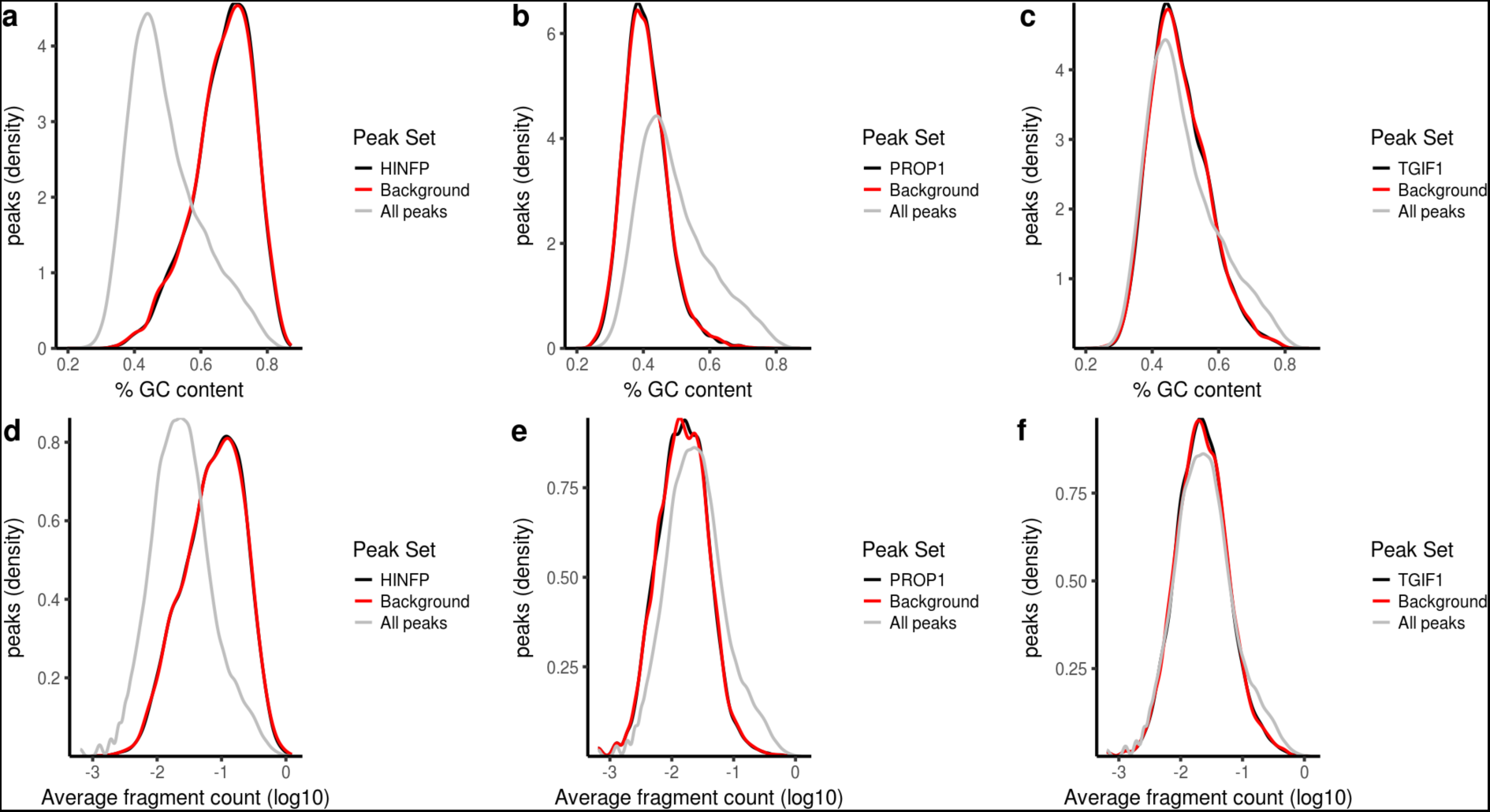
Examples of background matched peak sets. (a-c) Distribution of GC content per peak for all peaks (grey), the given motif of interest (black), and a background set for that motif (red). (b-f) Distribution of average fragment count per peak (log_10_) for all peaks (grey), the given motif of interest (black), and a background set for that motif (red). The distributions for GC content or average fragment count per peak for the background peak sets match very closely the distributions for the actual motif peak sets, even when the distribution for the motif peak set is quite distinct from that of all peaks.

**Figure S5.**
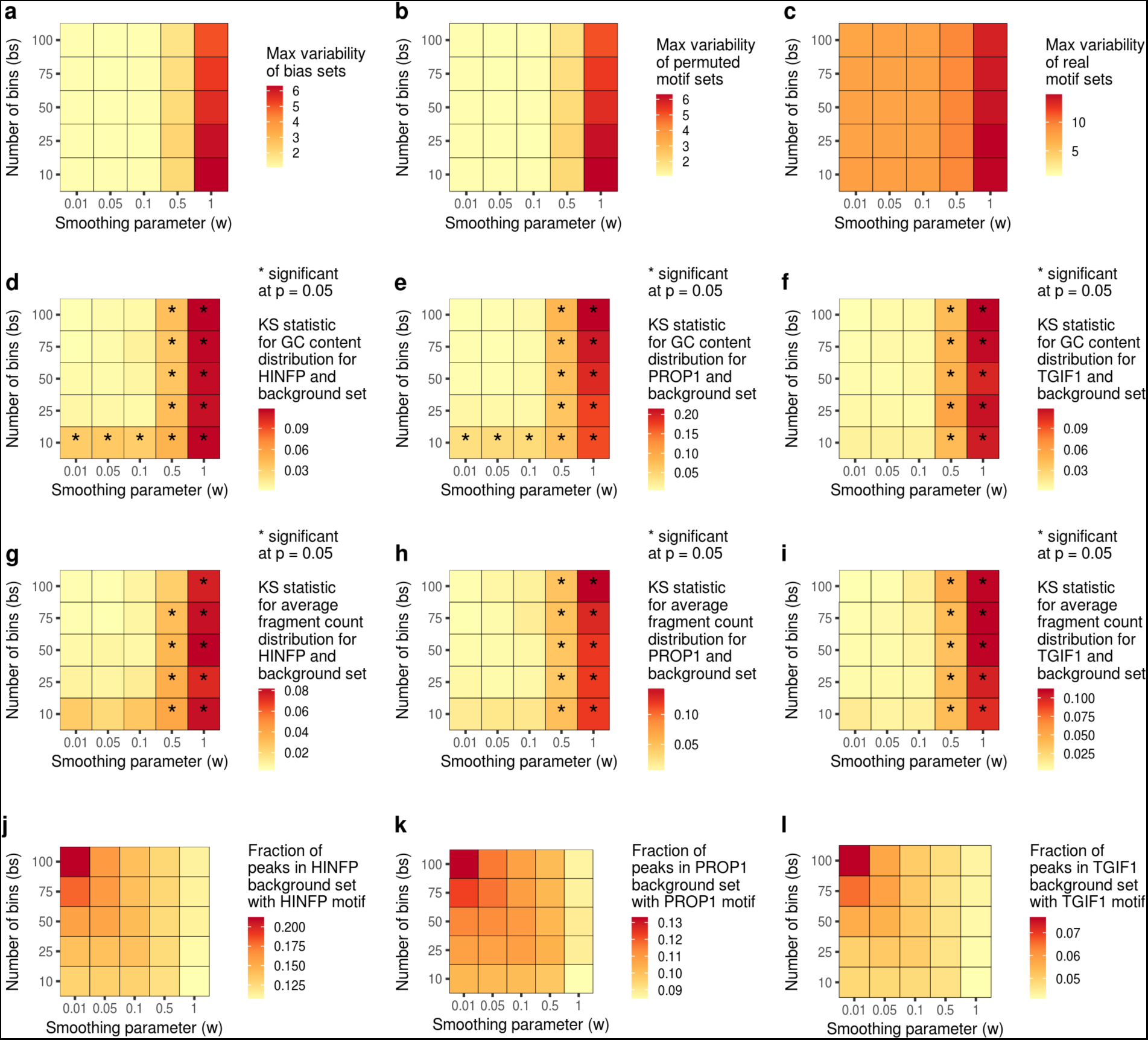
Effects of modifying parameters in background peak set selection. For each panel, the value of a different metric is shown based on varying the two parameters involved in background peak selection: (Rows) The number of bins (bs) used for grouping peaks based on GC content and average accessibility. This number represents the number of bins along one dimension; the actual number of bins is the square of this value. (Columns) A smoothing parameter (w) that controls how likely a peak in a bin will be chosen as a background peak for a peak in another bin. a) The maximum variability of a collection of peak sets selected to be constrained in GC content, average accessibility, or both. b) The maximum variability of a collection of peak sets selected to match the GC content & average accessibility of JASPAR motif peak sets c) The maximum variability for the JASPAR motifs. d-e) The KS statistic for the distribution of GC content for three motifs and for one background peak set for each motif. g-i) The KS statistic for the distribution of average fragment count for three motifs and for one background peak set for each motif. j-l) The degree of overlap between the motif peak set and the background peak set for three motifs. This analysis of the parameter space suggests that our chosen default values for w (0.1) and bs (50) enable optimal identification of true variability with minimal false discovery due to confounding technical biases. With lower values of the bs parameter or higher values of the w parameter, the background sets do not match the motif sets as well in terms of the GC content and average accessibility, yielding high variability for random sets of peaks with skewed GC content and/or average accessibility. However, with higher values of the bs parameter or lower values of the w parameter, the background sets are more likely to contain large fractions of the target motif peak set itself, reducing power for capturing the variability of that motif peak set.

**Figure S6.**
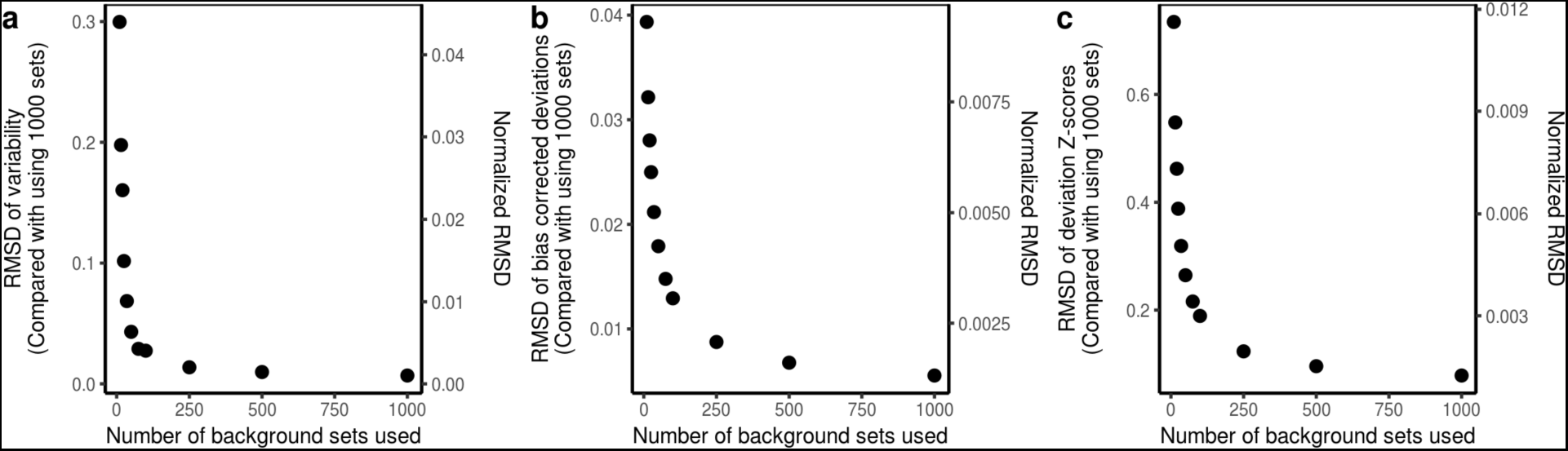
Differences in variability, bias corrected deviations, and deviation Z-scores based on number of background sets used. a) RMSD and normalized RMSD of variability when using a given number of background sets relative to when using 1000 background sets. Normalized RMSD is RMSD divided by the range of the variability when using 1000 background sets. b) Same as (a) but for bias corrected deviations instead of variability. c) Same as (a) but for deviation Z-scores instead of variability.

**Figure S7.**
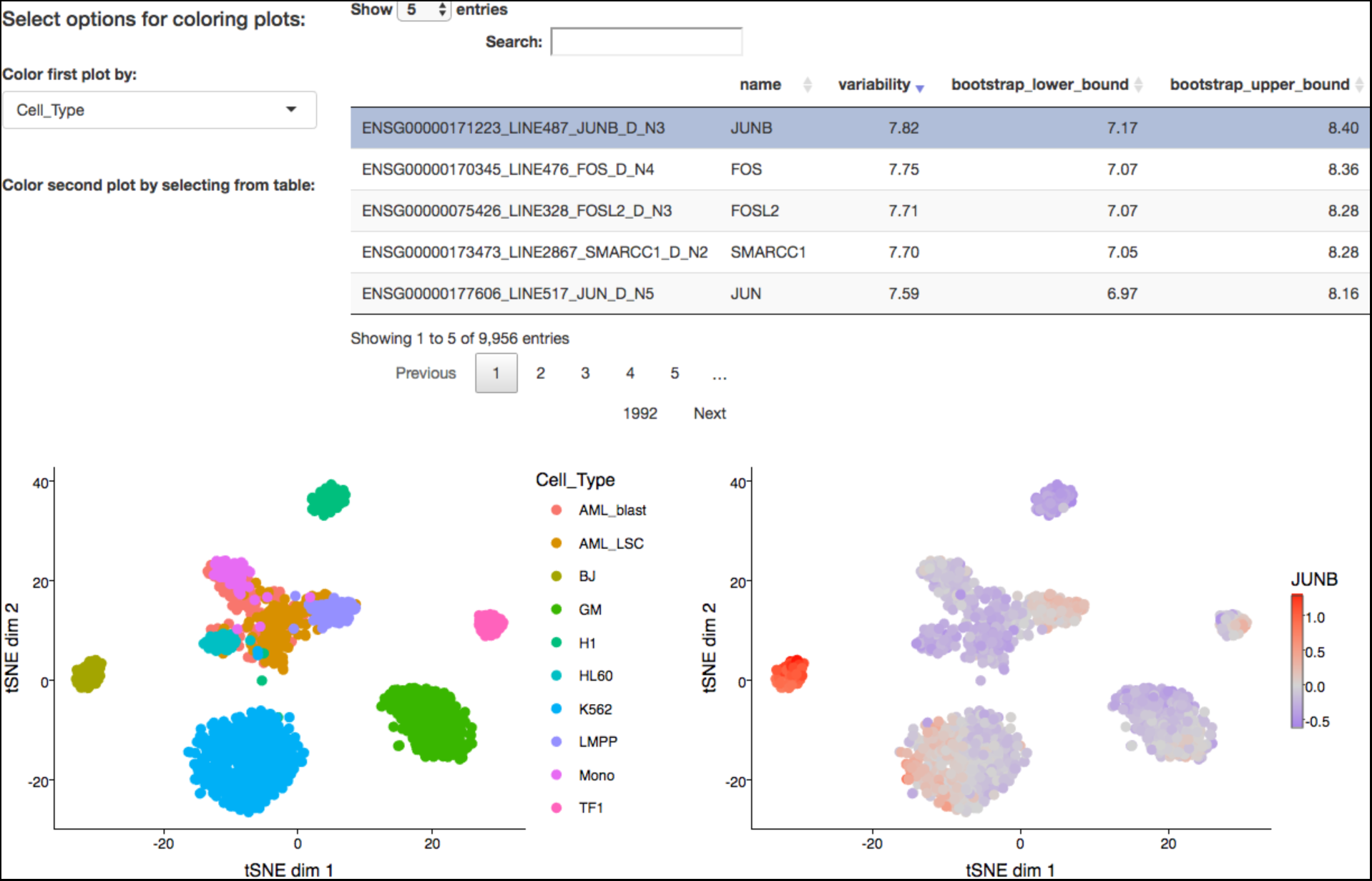
Screenshot example of interactive data browsing platform enabled by chromVAR. The plot on the left can be colored based on an annotation chosen from the dropdown menu above it. The plot on the right can be colored based on the bias corrected deviations of a motif chosen from the table above it. The table may be sorted based on any of the columns or searched for a particular name using the search box. The chromVAR R package includes a function for generating this type of interactive application.

**Figure S8.**
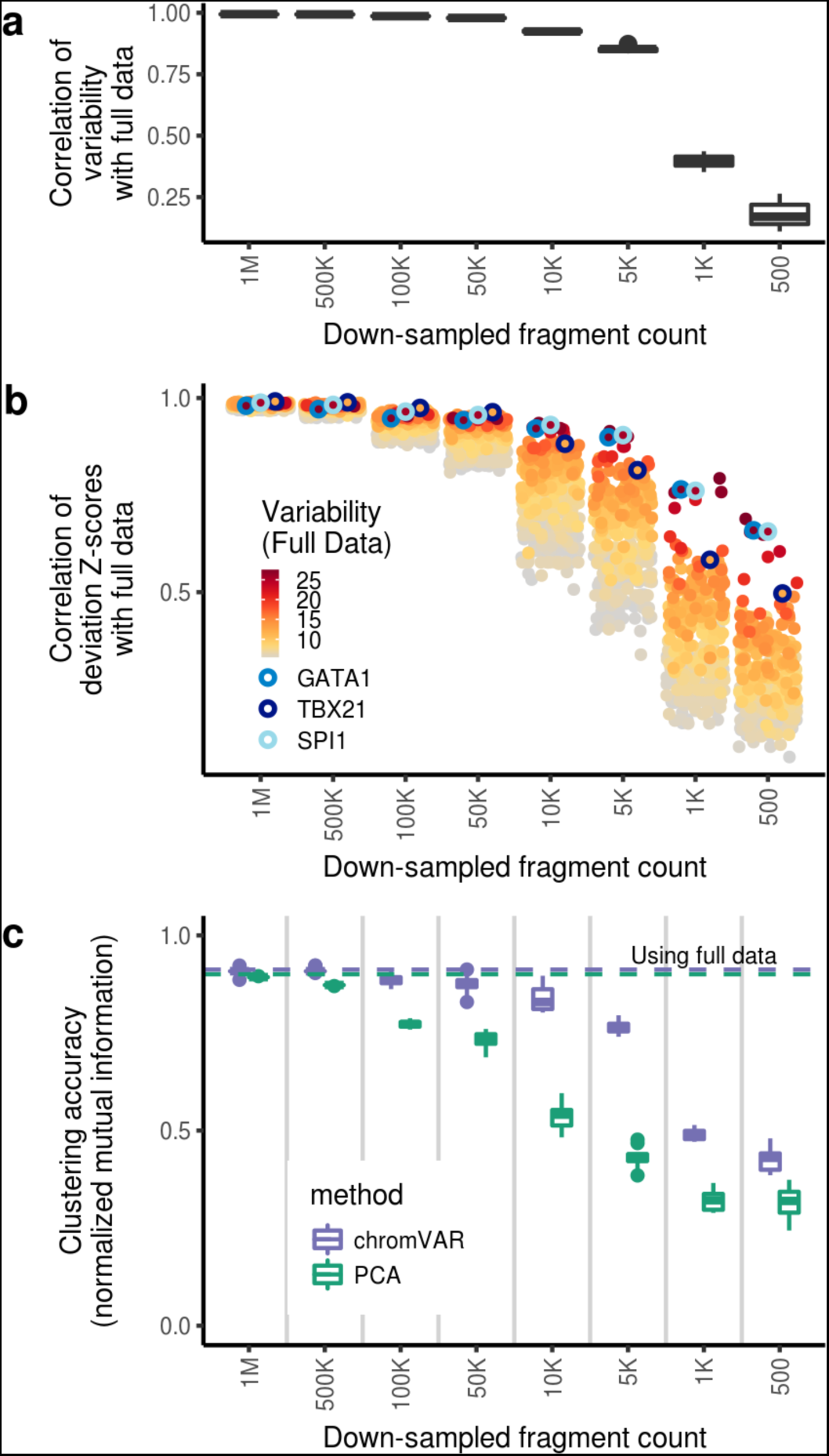
chromVAR results are robust to down-sampling and compare favorably to PCA. a) Correlation between chromVAR computed variability for down-sampled data versus full data set. Box plot shows results from 10 different down-sampling iterations. b) Correlation between chromVAR deviation Z-scores for down-sampled data versus full data set. Each point represents the median correlation for a different motif for the 10 downsampling iterations; only motifs in the top 20% of variability in the full data set are shown. Several key regulators are highlighted. c) Clustering accuracy when clustering based on chromVAR bias corrected deviations or distance between the top five principal components after down-sampling.

**Figure S9.**
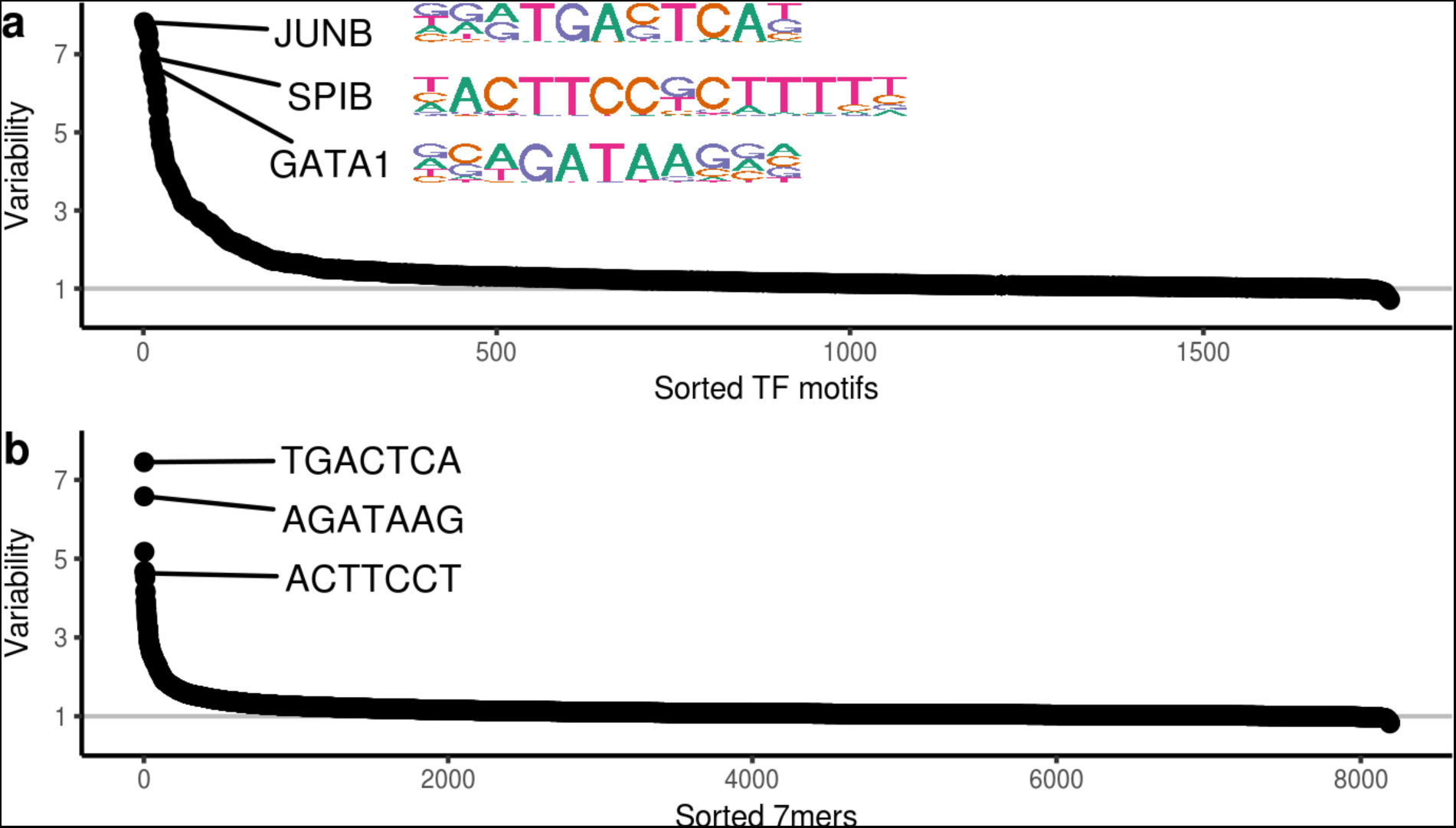
chromVAR variability. Variability in chromatin accessibility across cells for peak sets defined by a) motifs or b) 7mers.

**Figure S10.**
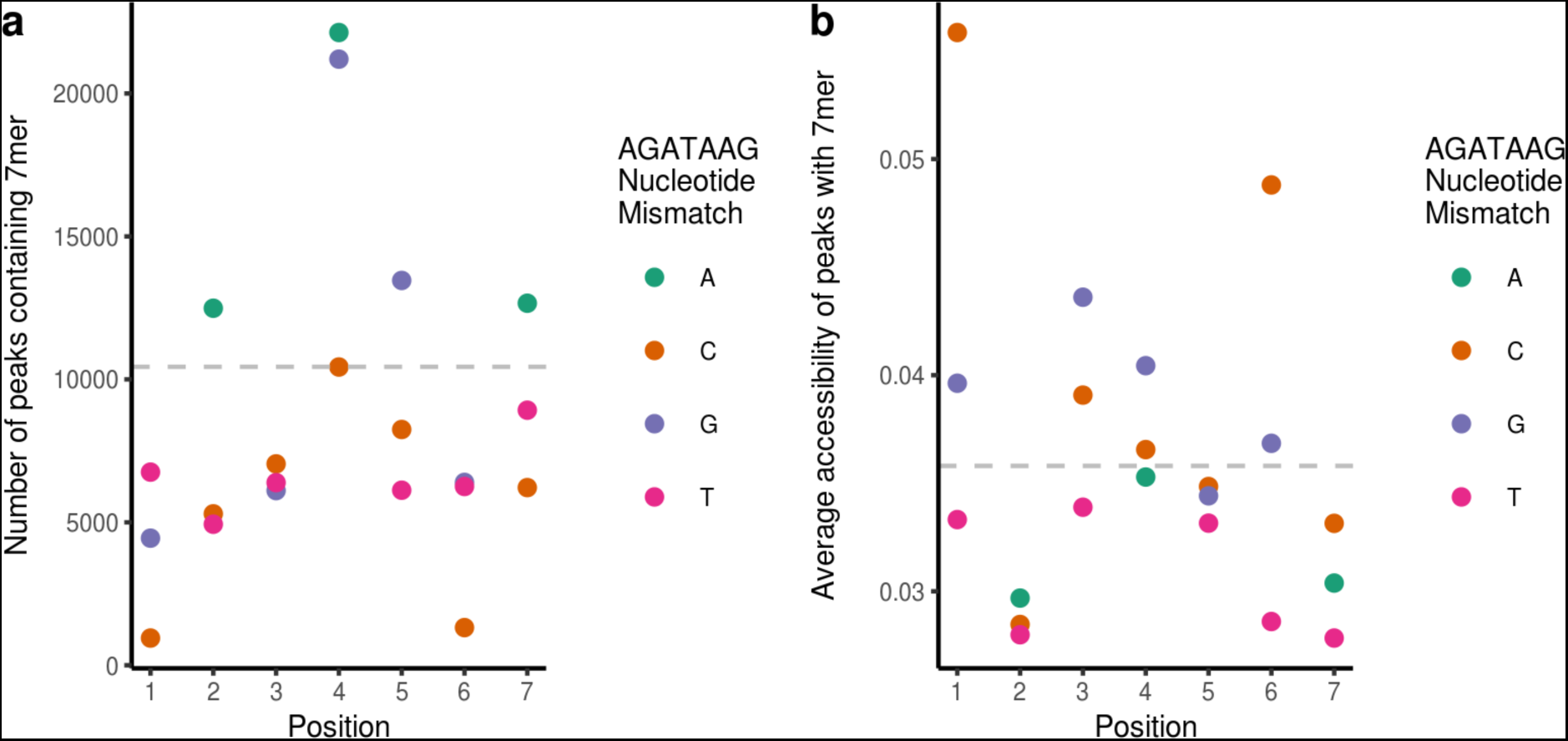
Number and accessibility of peaks containing kmers with 1 mismatch from AGATAAG. The gray dotted line indicates the number of peaks (a) or average accessibility of peaks (b) with the AGATAAG kmer.

**Figure S11.**
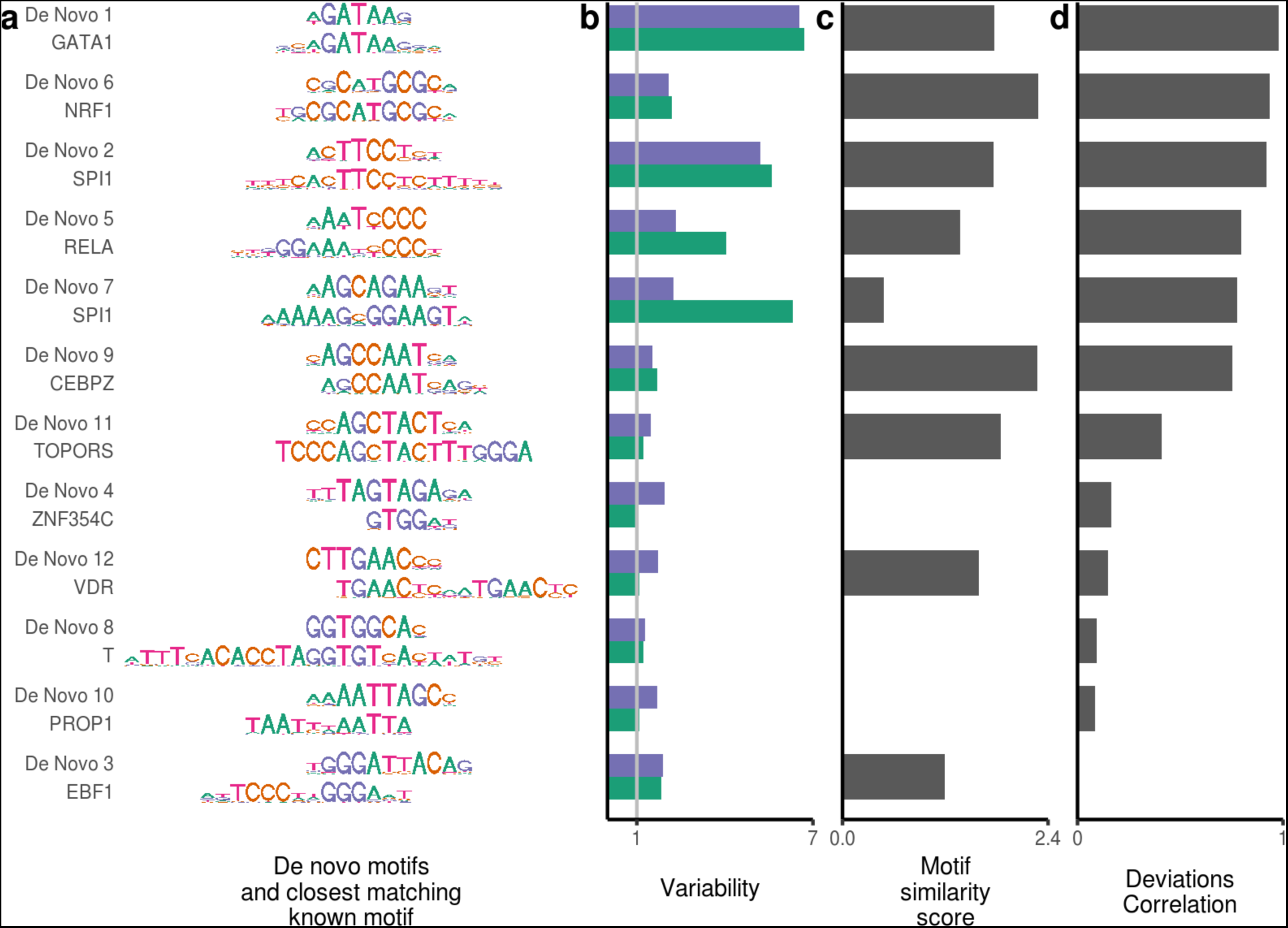
Motifs identified *de novo* by chromVAR from accessibility deviations for 7mers. (a) Each *de novo* motif is shown with the closest matching known motif below it. (b) The variability for both the *de novo* motif and the known motif (c) motif similarity score between the *de novo* motif and the known motif (see methods). (d) the correlation between the bias corrected deviations of the *de novo* motif and the known motif.

**Figure S12.**
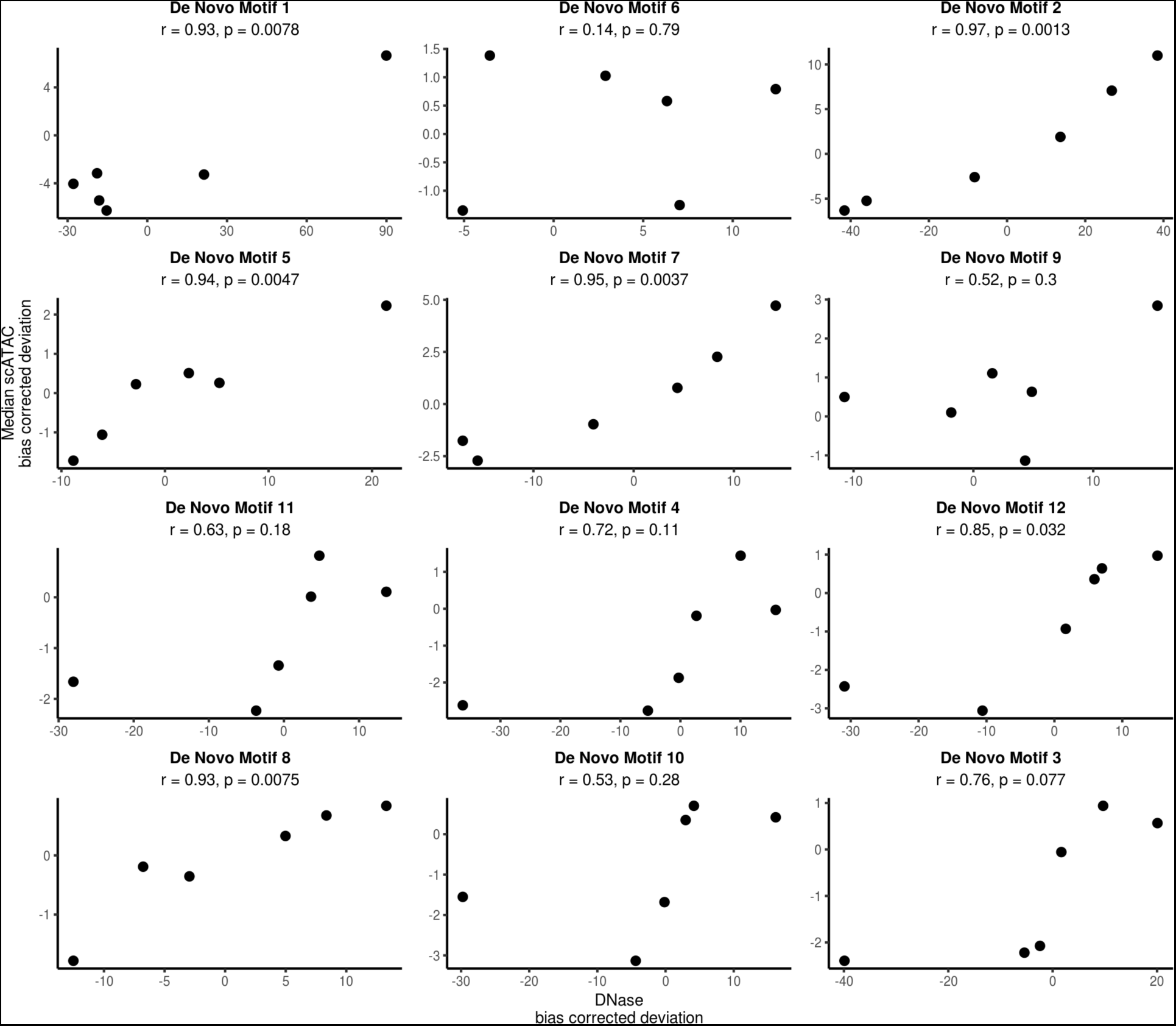
Comparison of accessibility variation between single cell ATAC-seq and DNase data for de novo motifs identified by chromVAR. Each panel corresponds to a different *de novo* motif and shows the median DNase deviation Z-score along the x-axis and the median scATAC Z-score along the y-axis. See Table S1 for mapping between samples:

**Table S1.** Mapping between Roadmap DNase samples and scATAC for Figure S12.

| scATAC-seq cell type | Roadmap DNase cell type |
| --- | --- |
| H1 | H1 Cell Line and H9 Cell Line |
| Monocytes | Primary monocytes from peripheral blood and Monocytes-CD14+ RO01746 Cell Line |
| GM12878 | GM12878 Lymphoblastoid Cell Line |
| K562 | K562 Leukemia Cell Line |
| BJ | Foreskin Fibroblast Primary Cells skin01 and Foreskin Fibroblast Primary Cells skin01 |
| LMPP | Primary hematopoietic stem cells G-CSF-mobilized Female and Primary hematopoietic stem cells G-CSF-mobilized Male |

**Figure S13.**
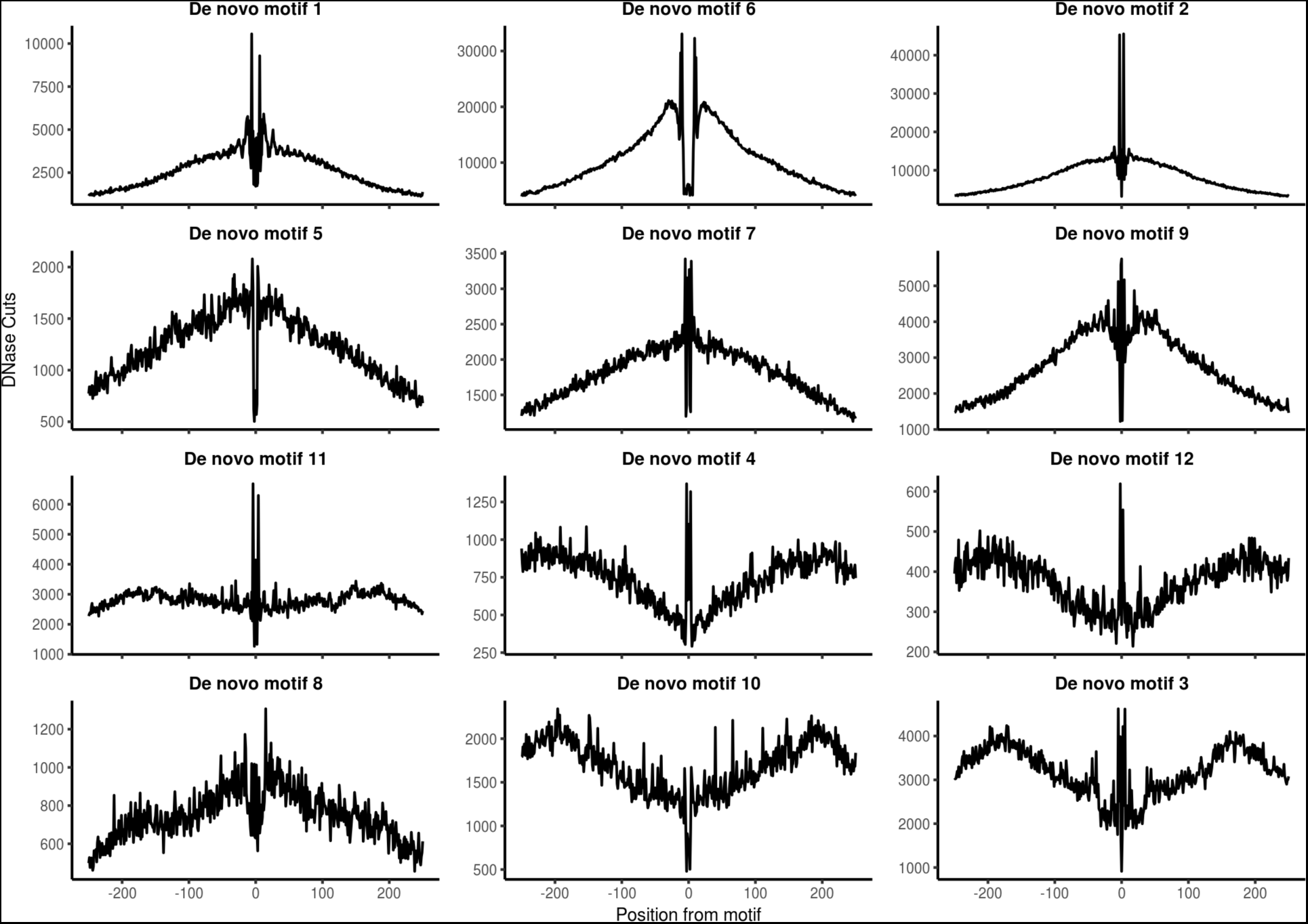
DNase footprints at *de novo* motifs identified by chromVAR. Each panel shows the DNase profile aggregated around all *de novo* motifs in the DNase sample with the highest accessibility deviation.

